# LRH-1/NR5A2 regulates the PTGS2-PGE_2_-PTGER1 signalling axis contributing to islet survival and antidiabetic actions of the agonist BL001

**DOI:** 10.1101/2021.11.02.466895

**Authors:** Eugenia Martin Vázquez, Nadia Cobo-Vuilleumier, Raquel Araujo Legido, Emanuele Nola, Lucia López Bermudo, Alejandra Crespo, Silvana Y. Romero-Zerbo, Maria García-Fernández, Alejandro Martin Montalvo, Anabel Rojas, Valentine Comaills, Francisco J. Bérmudez-Silva, Maureen Gannon, Franz Martin, Petra I. Lorenzo, Benoit R. Gauthier

## Abstract

We have previously described a role of LRH-1/NR5A2 in islet morphogenesis during postnatal development and reported that the treatment with BL001, an agonist of LRH-1/NR5A2, protects islets against-stress induced apoptosis and reverts hyperglycemia in 3 mouse models of Type 1 Diabetes Mellitus (T1DM). Islet transcriptome profiling revealed that most differentially expressed genes after BL001 treatment are involved in immunomodulation, among them, the increase in PTGS2/COX2 expression. Herein, we dissected the cellular and molecular branches of the BL001/LRH-1/NR5A2 signalling axis in order to chart the mode of action confering beta cell protection and hyperglycaemia reversion. We found that constitutive LRH-1/NR5A2 ablation within the insulin expression domain (RIP-Cre mouse model) caused a significant beta cell mass reduction characterized by blunted proliferation correlating with animal growth retardation, weight loss and hypoglycemia, leading to lethality before weaning. Using an inducible approach (pdx1^PB^CreER^™^ mouse model), specific deletion of LRH-1/NR5A2 in adult beta cells abolished the anti diabetic effect of BL001 in streptozotocin treated mice, correlating with complete beta-cell mass destruction. Additionally, BL001 induced *Ptgs2* expression, was blunted in islets lacking LRH-1/NR5A2. The combined BL001/cytokine treatment did not further stimulate *Ptgs2* expression above levels detected with cytokine alone yet secreted PGE_2_ levels were increased 5-fold. Inactivation of PTGS2 blunted induction of the target and its product PGE_2_ in islets treated with cytokines alone or with BL001. Importantly, PTGS2 inactivated islets were refractory to the BL001 protective effect under cytokine attack as evidenced by increased *Bax* expression levels, cytochrome C release and cleaved PARP. The PTGER1 antagonist ONO-8130, but not the PTGER4 antagonist L-161,982, negated BL001-mediated islet survival. Our results establish that the beneficial properties of BL001 against stress-induced cell death are specifically conveyed by LRH-1/NR5A2 activation in beta cells and downstream stimulation of the PTGS2-PGE_2_/PTGER1 signalling axis.

## INTRODUCTION

Nuclear receptors (NRs) are key factors of a vast number of physiological and pathophysiological processes. At the cellular level, they control energy homeostasis, survival, and plasticity via genetic and epigenetic regulation. At the whole organism level NRs participate in development, reproduction, metabolism, immune response and tissue regeneration ^1^. Ligand dependent regulation of NRs activity has forged them as druggable factors with promising therapeutic value for cancerous, metabolic and immune diseases. Of particular interest, is the liver receptor homolog 1 (LRH-1, also known as NR5A2) that belongs to the NR5A family of nuclear receptors ^2^. LRH-1/NR5A2 is highly expressed in the liver, intestine and pancreas, whereas lower expression levels are detected in the brain and immune cell types such as macrophages and T cells ^3^. LRH-1/NR5A2 is essential for embryonic development, while in adult it regulates key metabolic pathways. In the liver LRH-1/NR5A2 modulates the expression of target genes involved in glucose, cholesterol and bile acid metabolism, attenuates the hepatic acute phase response triggered by pro-inflammatory cytokines, and protects against endoplasmic reticulum stress ^4^. In the intestine, LRH-1/NR5A2 modulates the enterocyte renewal and regulates the local immune system via production of glucocorticoids ^5^. In the pancreas, LRH-1/NR5A2 regulates the expression of genes involved in digestive functions, and protects the endocrine islets against cytokine- and streptozotocin-induced apoptosis ^6, 7^. Haploinsufficiency of NR5A2 sensitizes mice to pancreatic cancer highlighting the role of NR5A2/LRH1 as a pancreatic tumour suppressor rather than an oncogenic factor ^8, 9^.

Given the role of LRH-1/NR5A2 in beta cell survival, glucose homeostasis and in attenuating inflammatory processes, attempts to regulate its activity is of important therapeutic value for the treatment of diabetes. Accordingly, administration of dilauroyl phosphatidylcholine (DLPC, the natural ligand of LRH-1/NR5A2) was shown to decrease hepatic steatosis and improve glucose homeostasis in two mouse models of insulin resistance and T2DM ^10^. These results prompted the development of hybrid phospholipid mimics as well as small non-polar bicyclic compounds that act as potent LRH-1/NR5A2 ligands ^11, 12, 13^. Based on one of these published structures able to bind to the ligand binding pocket of NR5A2/LRH1, we synthesized a small chemical agonist of LRH-1/NR5A2, denoted as BL001, and assessed its anti-diabetic therapeutic value for T1DM. We found that BL001 treatment either prophylactically or therapeutically reduced the incidence of diabetes in 3 independent pre-clinical mouse models of T1DM and protected human islets against stress-induced apoptosis ^14, 15^. These anti-diabetic benefits of BL001 were conveyed via immune cell tolerization and regression of insulitis combined with enhanced islet beta cell expansion and survival, a process that we have coined as ‘immune coupled trans-regeneration^16^. Such coupling was further substantiated by the finding that most of differentially regulated genes in BL001-treated mouse islets were involved in immunomodulation indicating a dialogue between islet cells and immune cells ^14^. Among these target genes stimulated by BL001 was the inducible prostaglandin endoperoxide synthase-2 (PTGS2 a.k.a. COX-2) that has been previously shown to partially protect mouse islets from streptozotocin (STZ)-mediated diabetogenic toxicity as well as to enhance human islet survival in the presence of cytokines through interaction with one or multiple prostaglandin E2 receptors (PTGERs) ^17, 18^. Notwithstanding, other studies contend that inhibition or deletion of PTGS2 shield human islet against cytokine-mediated beta cell dismay ^19^. Interestingly, in breast cancer cells the downstream metabolic pathway product of PTGS2/COX2, prostaglandin E2 (PGE_2_), induces LRH-1 mRNA expression by recruitment of multiple transcriptional activators to the LRH-1 promoter ^3^. Taking all together, whether the axis LRH1/PTGS2 is of importance in pancreatic beta cells remains to be clarified.

LRH-1/NR5A2 ablation in beta cells using the constitutive RIP-Cre/LRH-1^lox/lox^ mouse model resulted in early lethality of the mice (before weaning) ^14^, hampering the attempt to demonstrate both the specificity of LRH-1/NR5A2 to convey BL001-mediated islet cell survival and more importantly to delineate its mode of action. Analysis of the pancreatic islets of these mice at P1 (post-natal day 1) revealed a reduced number of pancreatic beta cells indicating that LRH-1/NR5A2 deletion at the onset of beta cell identity commitment likely perturbs beta cell function or mass^14^. Herein, we sought to: 1) further delineate the role of LRH-1/NR5A2 in islet morphology and architecture postnatally, 2) determine whether the anti-apoptotic and anti-diabetic effects of BL001 are specifically conveyed via LRH-1/NR5A2 activation in beta cells and 3) delineate the BL001/LRH-1/NR5A2 anti-apoptotic mode of action in beta cells with an emphasis on the potential implication of the PTGS2 signaling cascade ^16^. We report that deletion of LRH-1/NR5A2 blunts neonatal beta cell proliferation consistent with growth retardation, weight loss and hypoglycemia. Noteworthy, Cre expression and thus LRH-1/NR5A2 deletion was also detected in brain suggesting a potential contribution of this organ to the severe phenotype. We further report that LRH-1/NR5A2 deletion in adult beta cells abolishes the antidiabetic properties of BL001 an effect that involves the pro-survival PTGS2/PGE_2_/PTGER1 signaling axis.

## Materials and Methods

### Mice and regimen

All experimental mouse procedures were approved by the Institutional Animal Care Committee of the Andalusian Center of Molecular Biology and Regenerative Medicine (CABIMER) and performed according to the Spanish law on animal use RD 53/2013. Animal studies were performed in compliance with the ARRIVE guidelines ^20^. C57BL/6J mice were purchased either from Charles Rivers (L’Arbresle Cedex, France) or Janvier Labs (Saint-Berthevin Cedex, France). Two triple transgenic mouse models were generated: 1) a constitutive and beta cell specific LRH-1/NR5A2 ablation mouse model, in which the LRH-1^lox/lox^ mouse line was bred to the hemizygous RIP-Cre mouse line that constitutively express the Cre recombinase under the control of the rat insulin promoter (RIP) and 2) an inducible cell specific LRH-1/NR5A2 ablation mouse model, in which the LRH-1^lox/lox^ mouse line was bred to the pdx1^PB^CreER^™^ mouse line that harbours a tamoxifen (TAM)-inducible Cre recombinase/estrogen receptor fusion protein under the transcriptional control of the PDX1 promoter ^21^. These two double transgenic mouse lines were then bred to the Rosa26-YFP reporter mouse line to generate the ConβLRH-1KO and IndβLRH-1KO mouse lines respectively. Mice were housed in ventilated plastic cages under a 12-h light/dark cycle and given food and water *ad libitum*. For experiments conducted with ConβLRH-1 mice, P1, P7, P14 and P21 pups generated from wild type (LRH-1^+/+^), heterozygous (LRH-1^-/+^) and null (LRH-1^-/-^) were euthanized and pancreas extracted to perform immunofluorescence (IF) and morphometric analyses. Of note, as aging RIP-Cre mice may display beta cell impairment, control WT pups included RIP-Cre as well as *Lrh1/Nr5a2*^lox/lox^ pups ^22^. For experiments using IndβLRH-1KO mice, TAM (Sigma-Aldrich, Madrid, Spain, 20 mg/ml dissolved in corn oil) was administered via gavage for 5 consecutive days to 8-week old homozygous IndβLRH-1^-/-^ male or female mice (0.2 mg TAM/g body weight). This age was selected in order to have beta cell specific deletion of LRH-1/NR5A2 in adult mice. Animals were allowed a 4 weeks recovery period in order to eliminate any trace of TAM that may impact metabolism ^23, 24^. Mice were then intraperitoneally (*i.p*.) injected with 10 mg/kg body weight BL001/Vehicle daily for 5 weeks as previously described ^14^. BL001 treatment was initiated 1 week prior to an *i.p*. injection of a single high dose of STZ (175 mg/kg body weight). Circulating glucose levels were measured from venepuncture-extracted blood samples using a Precision Xceed glucometer (Abbott Scientifica SA, Barcelona, Spain). Mice with blood glucose above 250mg/dL for two consecutive measurements were considered hyperglycaemic. For Oral Glucose Tolerance Tests (OGTTs), mice were fasted for 5 hours and then received a glucose bolus (3g/kg body weight) by gavage. Blood was taken by venepuncture and circulating glucose was determined at different time points. At termination of experiments, mice were euthanized and organs (pancreas, liver and brain) extracted for further analysis (QT-PCR and immunofluorescence). For transaminases analysis, blood was collected from the tail vein in a sterile environment. Liver damage was induced using diquat dibromide (i.p. 125 mg/kg body weight). The analysis was performed using a Cobas Mira autoanalyzer and reagents from Spinreact (Spinreact S.A.U., Girona, Spain) and Biosystems (Biosystems S.A., Barcelona, Spain).

### Islet isolation and cell culture

Islets were isolated from either C57BL/6J mice or TAM and vehicle-treated IndβLRH-1KO mice and cultured as previously described with some modifications ^14^. Briefly, the isolation was performed by infusing the pancreatic duct with collagenase followed by pancreas digestion, and islet purification. For the infusion, the ligation of the common bile duct in close proximity to the pancreas was executed with one suture. The duodenum wall, near to the edge of the ampulla, was then clamped both sides. An incision into the ampulla was performed with a 30-G needle. Two to three ml of collagenase V solution (Sigma, 1mg/ml) was infused trough the incision with a blunt 30-G needle coupled to a 3 ml syringe. Pancreas was then removed and digested at 37°C for 5+5 minutes shaking 20 times in between the two times. Islets were then handpicked. The RAW264.7 mouse macrophage cell line (ATCC, USA) was cultured in high glucose DMEM supplemented with 10% FBS and 2 mM L-glutamine, 100 U/ml penicillin and100 mg/ml streptomycin.

### RNA interference

Islets were transfected with either 50 μMol of a pool of *PTGS2* small interfering (si)RNAs ON-Target plus Mouse Ptgs2 (19225) siRNA-SMARTpool (Dharmacon/Horizon Discovery, Cultek, Madrid, ES) or a scramble siRNA (Sigma-Aldrich) as previously described ^25^. Transfections were performed using the Viromer Blue siRNA/miRNA transfection kit following the manufacturers’ instructions (Origene, Rockville, USA). Forty eight hours post siRNA treatment, islets were treated or not with cytokines and/or BL001. Twenty four hours later islets were processed for RNA isolation, protein extraction or IF analysis. Media was also collected to measure PGE_2_.

### *In vitro* cell treatments

Islets were cultured in the presence of: 1) either 1 or 10 μM BL001 (optimal doses previously used ^14^) and/or 2) a cocktail of cytokines (2 ng/ml IL1β, 28 ng/ml TNFα and 833 ng/ml IFNγ; ProSpec-Tany TechnoGene Ltd, Ness-Ziona, IL) and/or 3) 100 nM L-161,982 (Cayman Chemical Coorp, Ann Arbor, USA) and/or 4) 100 nM ONO-8130 (Tocris Bioscience, Bristol, UK). In the latter two conditions, islets were treated for 12 hours with the respective antagonists (L-161,982 and ONO-8130) followed by a second treatment with the antagonists along with cytokines. BL001/vehicle was added 30 minutes later and cells were further incubated for for 24 hours. RAW264.7 cells were treated, or not, with 1 μg/ml LPS for 2, 6 and 24 hours. Cells were then processed for ELISA, RNA isolation, protein extraction or immunofluorescence analysis. In some instances, the media was also collected for ELISA analysis.

### Cell death

Cell death (apoptosis) was measured using the Cell Death Detection ELISA kit (Roche Diagnostics, Madrid, Spain) as previously described ^26^.

### RNA extraction and quantitative real-time PCR

Total RNA from islets was extracted using the RNeasy Micro Kit (Qiagen) while RNA from liver and brain was extracted using TRIzol (Sigma-Aldrich). Complementary DNA using 0.1 to 1 μg RNA was synthesized using the Superscript III Reverse Transcriptase (Invitrogen-Thermo Fisher Scientific, Madrid, Spain). The QT-PCR was performed on individual cDNAs using SYBR green (Roche) ^14^. Gene-specific primers were designed using Primer3Web (https://primer3.ut.ee/) and the sequences are listed in **Table S1**. Expression levels were normalized to various reference genes including *Cyclophilin*, *Rsp9*, *Gapdh* and *Actin*. The relative gene expression was calculated using the standard curve-based method ^25^.

### Immunofluorescence analyses

For immunostaining, pancreas or brain were fixed overnight in 4% paraformaldehyde at 4°C. Pancreas were dehydrated, paraffin embedded, and sectioned at 5 μm thickness. Brains were coronally sectioned into six series of 50 μm slices using a vibratome (Leica). Isolated islets were processed and paraffin embedded as previously described ^27^. RAW264.7 cells were washed with PBS and fixed 15 minutes in 4% paraformaldehyde at room temperature. They were washed again with PBS and permeabilized with PBS-0,1% triton 15 minutes on ice. Tissue sections / cells / free floating brain sections were blocked in PBS containing 5% donkey/goat serum and 0.2%Triton X100 for 1h at R/T. Immunostaining was then performed overnight at 4°C using a combination of primary antibodies (Table S2). Subsequently, secondary antibodies were incubated for 1 hour at room temperature in PBS 0.2% TritonX100 (Table S3). Nuclei were stained with 0.0001% of 4’,6-diamidino-2-phenylindole (DAPI, Sigma-Aldrich) and cover slips were mounted using fluorescent mounting medium (DAKO). Epifluorescence microscopy images were acquired with a Leica DM6000B microscope and z-stack images were acquired using a Confocal Leica TCS SP5 (AOBS) microscope.

### Morphometric analyses

For the assessment of the whole pancreatic area, images of pancreatic sections were automatically acquired using the NIS-Elements imaging software. Morphometric quantification was performed using Photoshop, ImageJ and FIJI softwares. For ConsβLRH-1 mouse line, sections from 3 independent mice with an average of about 50 islets and 5000 cells per mice were used for quantifications. For IndβLRH-1-R26Y mouse line, sections from 3 to 6 independent animals with an average of about 50 islets and 4000 cells per group were used for quantifications.

### Western blot

Islets and RAW264.7 cells were processed and protein concentration determined as previously reported ^26^. Western blotting was performed as previously described ^28^. Antibodies and dilutions employed are provided in Table S2 and S3.

### Prostaglandin E_2_ (PGE_2_) ELISA

Media was collected from cultured islets after treatments, sieved through a 0.22 μm filter and diluted 3-fold in assay buffer. Secreted PGE_2_ levels were measured by ELISA, following the manufacturer’s instructions (R&D systems, Minneapolis, USA). Absorbance in each well was measured by using a Varioskan Flash Spectral Scanning Multimode (ThermoFisher Scientific) and serum PGE_2_ concentrations were calculated from the standard curve.

### Statistical analyses

Data are presented as means ± SEM. Student’s t-test or One-Way ANOVA were used as described in figure legends. *p* values less than or equal to 0.05 were considered statistically significant. Statistical analyses were performed using the GraphPad Prism software version 8 (GraphPad Software, La Jolla, USA).

## Results

### LRH-1/NR5A2 ablation disrupts neonatal beta cell replication

Constitutive deletion of the *Lrh1/Nr5a2* gene subsequent to beta cell commitment during development using the RIP-Cre::Lrh1^lox/lox^ mouse model results in offspring lethality between P1 and P21 ^14^, which was further evidenced by a decrease in homozygous knockout genotype frequency observed between P1 and P7 (**Figure 1a**). Prior morphometric analysis performed at P1 indicated that premature death may ensue from an inadequate postnatal islet beta cell pool ^14^, a premise we now extended to P14 and P21 in the RIP-Cre::Lrh1^lox/lox^::ROSA-YFP triple transgenic mouse model (ConβLRH-1KO mice) **(Figure 1b-e**). Although in the ConβLRH-1KO mice the number of total number of cells per islet as well as the number of alpha cells were decreased at P14 but normalized at P21 as compared to WT mice (litter match control mice with normal LRH-1/NR5A2 expression, including the RIP-Cre::ROSA-YFP and *Lrh1/Nr5a2*^lox/lox^::ROSA-YFP mice), the beta cell mass was significantly reduced at both time points in homozygous ConβLRH-1^-/-^KO-R26Y mice (**Figure 1b-e**). Interestingly, the percentage of YFP^+^/INS^+^ cells in homozygous ConβLRH-1^-/-^KO mice was significantly decreased as compared to heterozygous ConβLRH-1^-/+^ mice at both at P7 and P14 attaining approximately 60% of the overall beta cell population (**Figure 1f**). Although not significant, the percentage of YFP^+^/INS^+^ cells in heterozygous animals gradually decreased from 70% to 60% between P7 and P21 (**Figure 1f**). Active beta cell proliferation is a hallmark of neonatal islets that establishes the beta mass during adulthood both in humans and rodents ^29, 30, 31^. As such we reasoned that deletion of LRH1/NR5A2 may impact neonatal beta cell expansion. This premise was confirmed as evidenced by a drastic decrease in the percentage of Ki67^+^/INS^+^ cells in islets of homozygous ConβLRH-1^-/-^KO mice as compared to either heterozygous ConβLRH-1^-/+^or WT mice (**Figure 1g**). Consistent with reduced beta cell number Conβ LRH-1^-/-^KO mice displayed growth retardation correlating with an hypoglycemic state and weight loss as compared to either WT and heterozygous ConβLRH-1^-/+^ animals (**Figure 1h-j**). Noteworthy, the Cre transgene driven by the RIP was shown to be expressed also in the central nervous system (CNS) ^32, 33^. Consistent with these reports, we observed expression of YFP in several regions of the brain including the hypothalamus and cortex (**Figure S1a**). Since LRH-1/NR5A2 expression was also detected in the CNS, albeit at lower levels than in islets and liver (**Figure S1b**), we cannot exclude the involvement of neuronal LRH-1/NR5A2 ablation in the observed phenotype of ConβLRH-1^-/-^KO mice. Taken together these results demonstrate that LRH-1/NR5A2 contributes to neonatal beta cell expansion via cell replication.

**FIGURE 1:**
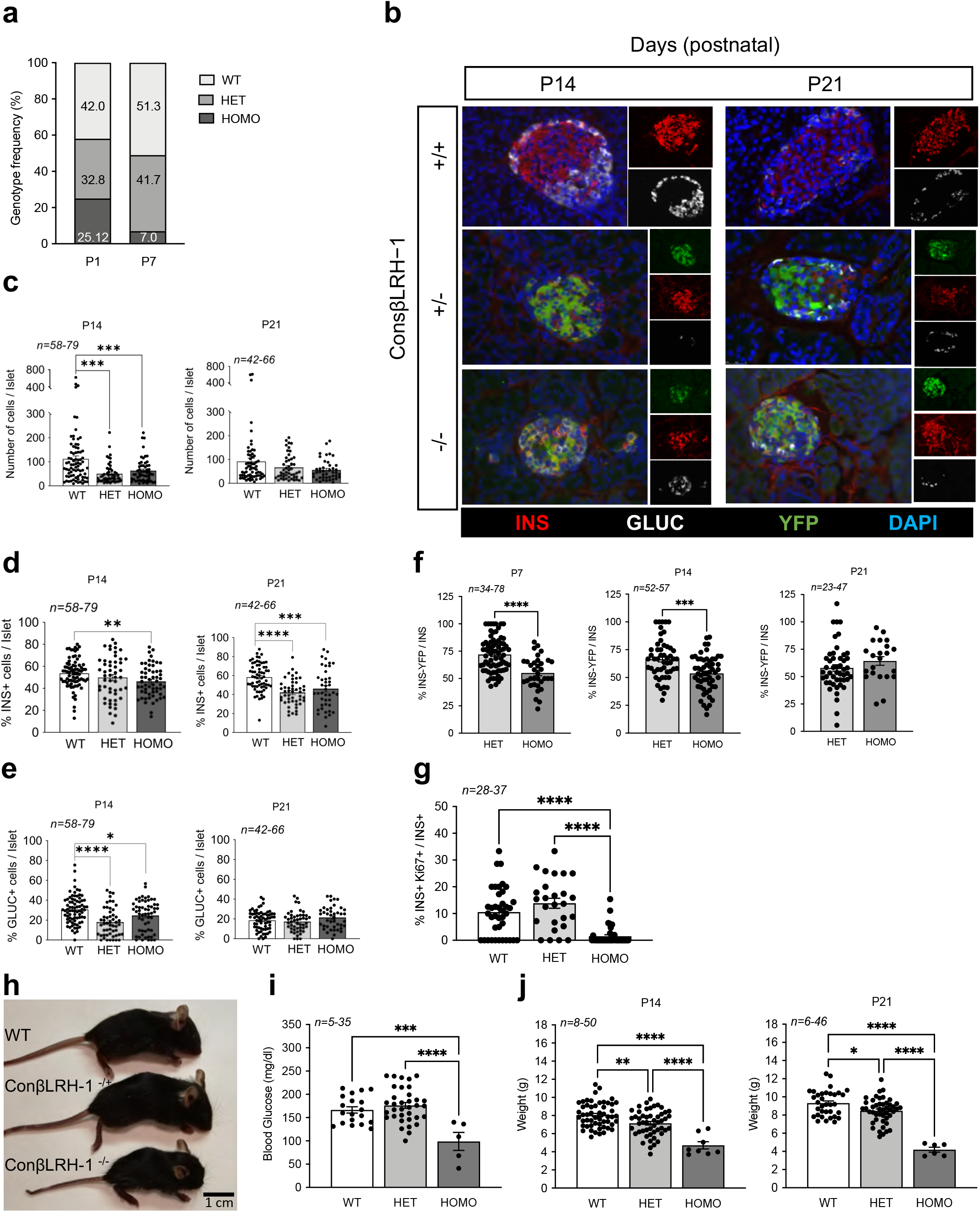
LRH-1 contributes to islet formation: (**a**) Representation of the genotypes frequency distribution: WT, LRH1wt/wt; HET, LRH1 wt/ko; HOMO, LRH1ko/ko). *n*=23 mice for P1 and n=347 mice for P7. (**b**) Pancreases were extracted at P14 and P21 and immunostaining for insulin (INS, red), glucagon (GLUC, white) and YFP (green) was performed on fixed sections. Morphometric analysis was performed to assess (**c**) islet cell composition, (**d**) β-cell mass (**e**) α-cell mass and (**f**) percentage of INS^+^/YFP^+^ cells. (**g**) Co-staining of Ki67 and Insulin was performed on P7 pancreas and percentage of double positives were assessed over total insulin positive cells (**h**) Representative images of wild type mice (WT) and ConβLRH-1 transgenic pups with either 1 (-/+) or 2 (-/-) *Nr5a2* allele disruption 21 days postnatal (P21). (**i**) Blood glucose levels (P21) and (**j**) weight of P14 and P21 WT, Hetero and Homozygous pups. * p<0.05, ** p<0.01, *** p<0.001 **** p<0.0001, one-way ANOVA, Tukey multiple comparisons test (**c,d,e, h** and **i**) and unpaired Student’s t test, HET versus HOMO (**f**).

### TAM-induced LRH1/NR5A2 deletion does not compromise the metabolic status of healthy IndβLRH-1KO mice

The goal of generating the ConβLRH-1KO transgenic mouse model was to demonstrate the specificity of BL001-mediated LRH-1 activation in conveying islet survival. However, as we uncovered that LRH-1/NR5A2 is important for postnatal islet beta cell mass expansion and its deletion compromised survival (**Figure 1**), we generated a second transgenic mouse model, the IndβLRH-1KO mice, in which LRH-1/NR5A2 is spatially and temporally deleted in adult beta cells in a TAM-dependent inducible manner using the PDX1-CreER^T^ mouse model. This deletion of LRH-1/NR5A2 in beta cells from adult animals avoids the developmental consequences of the lack of this nuclear receptor and minimizes off-target deletion ^21^. A 5-day TAM treatment did not alter body weight, glycemia nor glucose tolerance of IndβLRH-1KO mice either during or post-treatment (**Figure 2a-c**) independent of gender (**data not shown**). In addition, liver toxicity, measured by ALT and AST levels, was not observed at either 1 or 8 weeks post TAM-treatment (**Figure 2d-e**). As such all subsequent experiments were performed at 4 weeks after TAM treatment in order to ensure the elimination of any traces of the drug. TAM treatment of IndβLRH-1KO mice induced YFP expression in 70-80% of beta cells with a concomitant 80%reduction in NR5A2/LRH1 transcript levels in islet but not in the liver and more importantly not in the brain (**Figure 2f-i**). Altogether, these results indicate that TAM administration and beta cell specific ablation LRH-1/NR5A2 in adult beta cells do not compromise the metabolic health status of IndβLRH-1KO mice under normal physiological/environmental conditions.

**FIGURE 2:**
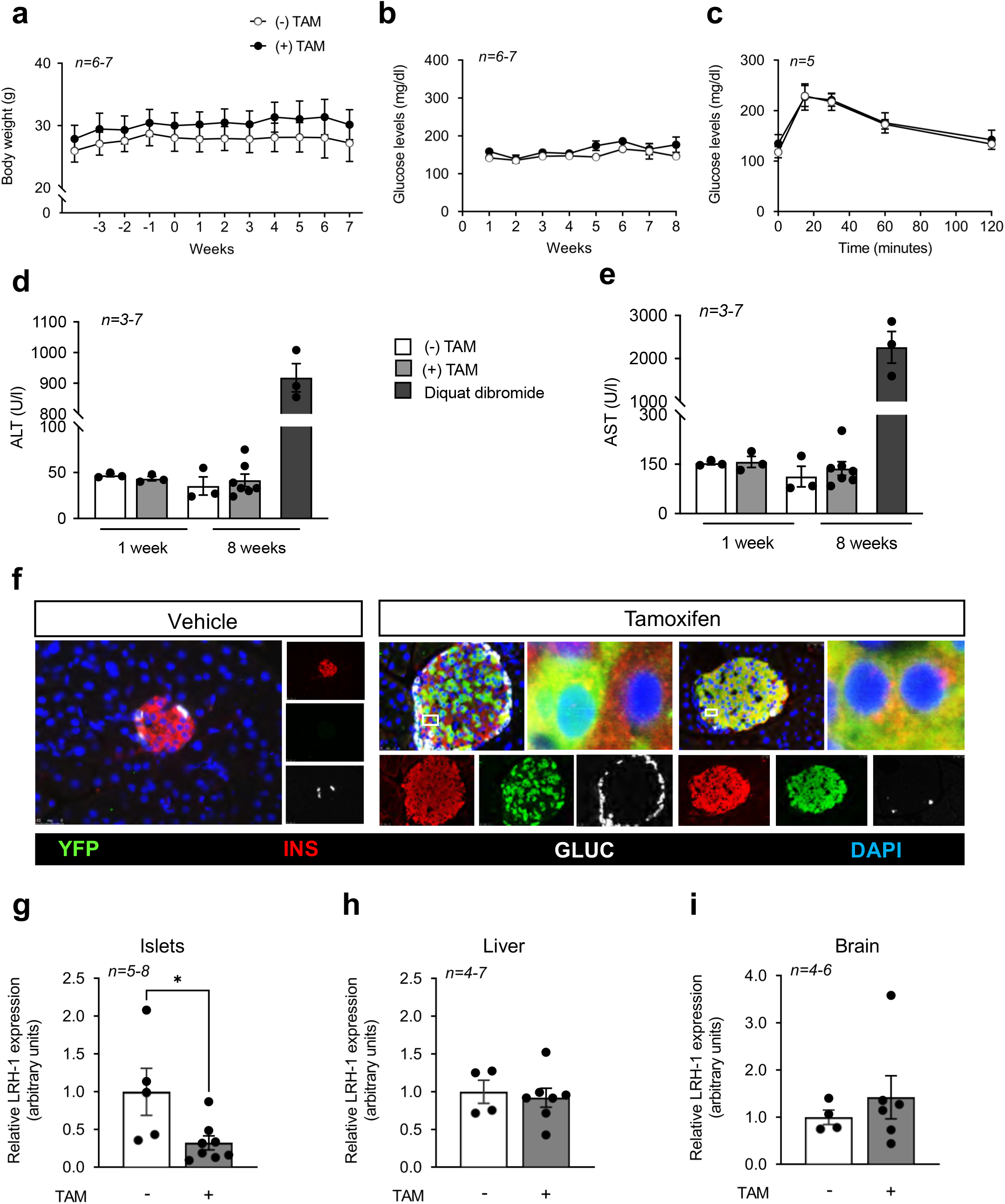
TAM treatment does not compromise the metabolic health status of 8 weeks old IndβLRH-1 mice. (**a**) The body weight of IndβLRH-1 treated or not with TAM was monitored for several weeks before (negative weeks) and after (positive weeks) treatment. TAM started at week 0 for 5 consecutive days (**b**) Blood glucose levels were also monitored starting at week 1 post TAM treatment. (**c**) An OGTT was conducted 4 weeks post TAM treatment. Circulating levels of (**d**) ALT and (**e**) AST were assessed 1 week and 8 weeks post TAM treatment. As positive control, liver damage was induced using diquat dibromide (125 mg/kg body weight). Results are expressed as means + s.e.m. (**f**) Representative images of pancreas sections from IndβLRH-1 treated or not with TAM co-stained for YFP (green), insulin (INS, red) and glucagon (GLUC, white). Nuclei were stained with DAPI. Magnification 40X. LRH-1 transcript levels were assessed in (**g**) islets, (**h**) liver and (**i**) whole brain extracted from IndβLRH-1 mice treated or not with TAM. Expression levels were normalized to the housekeeping gene *Cyclophilin* (CYCLO) or β-Actin. Results are expressed as means + s.e.m. *p < 0.05, unpaired Student’s t test, vehicle versus TAM.

### Conditional beta cell specific ablation of LRH-1/NR5A2 negates BL001-mediated cell survival and anti-diabetic properties

We next assessed the contribution of LRH-1/NR5A2 in mediating the beneficial antidiabetic and cell survival effects of BL001 in the IndβLRH-1 mice. As previously reported for WT mice after BL001 and STZ administration, only 20-30% of BL001 and non-TAM-treated IndβLRH-1 developed hyperglycaemia (**Figure 3a**). In contrast, after TAM treatment, and thus depletion LRH-1/NR5A2, 80% of mice developed hyperglycaemia subsequent to BL001/STZ treatment (**Figure 3b**). Therefore, the depletion of LRH-1/NR5A2 in beta cells blunts the anti-diabetic effect of BL001 treatment. Immunohistochemistry combined with morphometric analysis of the pancreas of these mice revealed that hyperglycaemic TAM/BL001/STZ-treated IndβLRH-1KO mice exhibited a drastic decrease in beta cell number, comparable to that of STZ treated mice (**Figure 3c and d**). In contrast normoglycemic non-TAM/BL001/STZ-treated IndβLRH1 mice harboured normal islet beta cell number similar to that of control non-STZ mice (**Figure 3c and d**). Interestingly, the number of alpha cells mass was increased in TAM BL001/STZ-treated mice as compared to non-TAM BL001/STZ-treated mice resulting in a decreased ratio of INS^+^/GCG^+^ cells (**Figure 3e and f**). These results establish that the anti-apoptotic and pro-survival properties of BL001 are specifically conveyed through its interaction with LRH-1/NR5A2 in beta cells.

**FIGURE 3:**
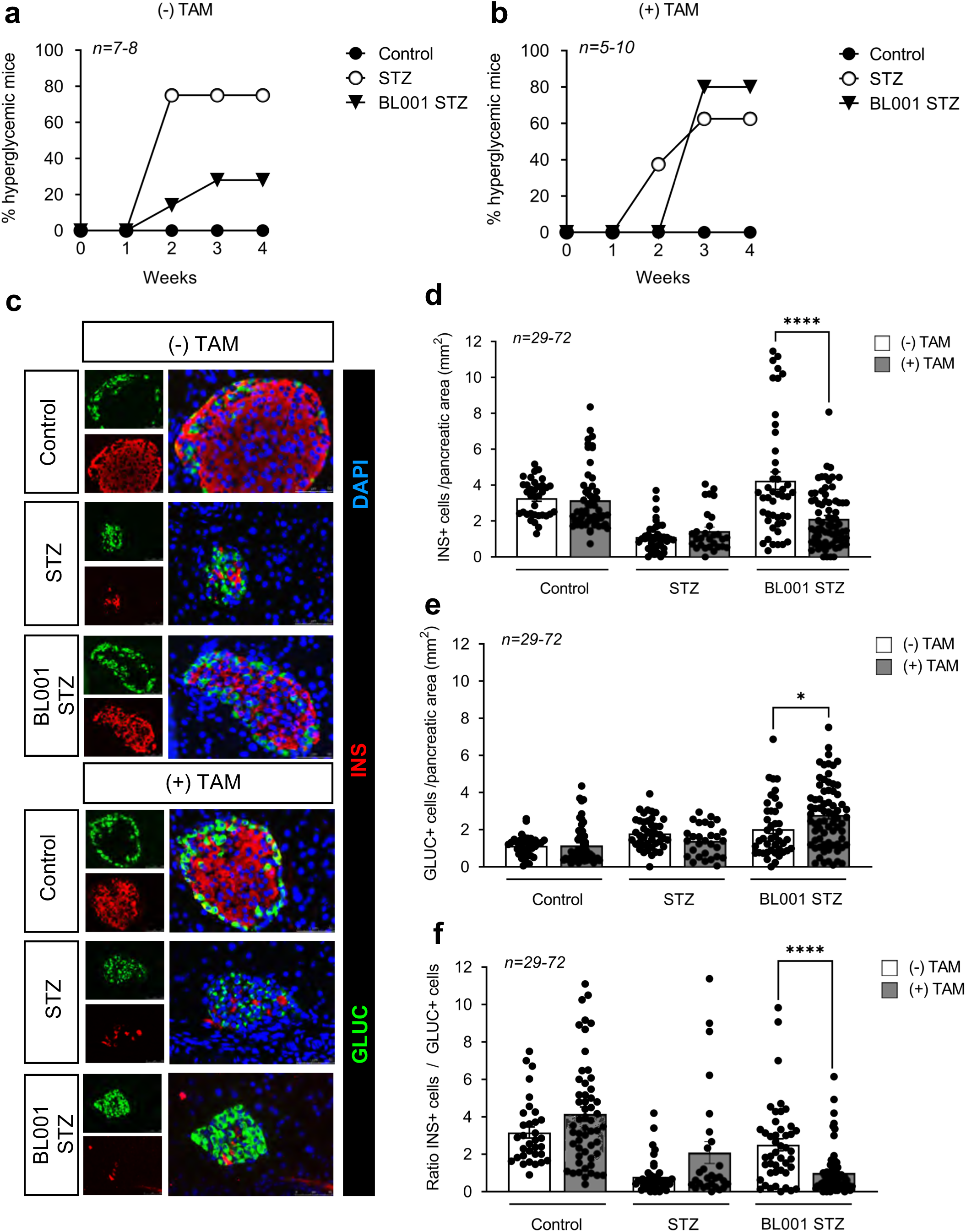
β-cell specific LRH-1 ablation in adult mice negates the anti-diabetic properties of BL001. Diabetes incidence of IndβLRH-1 mice either (**a**) not treated or (**b**) treated with TAM and then subjected to a STZ and/or BL001 regimen for 5 weeks. BL001 (10mg/kg body weight) treatment started 4 weeks after finishing TAM treatment. Diabetes was induced at week 1 post-BL001 treatment with a single high dose of STZ (175mg/kg body weight). (**c**) Representative immunofluorescence images of pancreas sections from the various experimental groups co-stained for insulin (INS, red) and glucacon (GLUC, green). Nuclei were stained with DAPI (blue). Magnification, 40X. Quantification of (**d**) β-cells, (**e**) α-cells and (**f**) ratio thereof in the different groups. n= individual islets counted from 3-6 independent mice. Data are represented as the mean ± s.e.m. *p < 0.05 and ****p < 0.0001, unpaired Student’s t test between (-) and (+) TAM provided in the various treatments.

### PTGS2 is a downstream target of both cytokines and the BL001/LRH-1/NR5A2 signalling pathway

Comparative global transcriptome profiling of BL001-treated and untreated mouse islets revealed that most of the differentially expressed genes are involved in immunomodulation pathways ^14^. Among these, is *Ptgs2* that generates PGE_2_ and that depending upon the levels of expression of the 4 different PGE_2_ receptors will convey either a pro- or anti-inflammatory stimuli ^34, 35^. To further dissect the molecular mechanism implicated in the BL001/LRH-1/NR5A2 signalling pathway conferring beta cell survival, we focused on PTGS2 as a downstream effector ^16^. Consistent with previous results, BL001 treatment increased *Ptgs2* expression, an effect that was abrogated in islets isolated from TAM-treated IndβLRH-1KO mice that lack LRH1/NR5A2, validating *Ptgs2* as a downstream target of LRH1/NR5A2 (**Figure 4a**). Islet *Ptgs2* expression was also increased by cytokines attaining a 3-fold stimulation at 24 h post treatment (**Figure 4b**). The latter time point was selected for subsequent experiments as most *in vitro* BL001 studies have been performed between 24 and 72 hours ^14^. Interestingly, the combined BL001/cytokine treatment did not further stimulate *Ptgs2* expression above levels detected with cytokine treatment alone (**Figure 4c**). Attempts were made, albeit unsuccessfully, to detect the PTGS2 protein in islets (**Figure S2a and b**). In contrast, PTGS2 was discerned in LPS-treated RAW264.7 cells correlating with a robust increase in its transcript levels suggesting that PTGS2 levels in islets are below the detection threshold (**Figure S2c-e**). To assess the contribution of PTGS2 in the BL001/LRH1 signalling pathway, the transcript was silenced using siRNA in isolated WT islets that were then challenged, or not, with a cytokine cocktail and further treated with BL001. Silencing of *Ptgs2* resulted in the dampened induction of the transcript by both cytokines and BL001 (**Figure 4d**). Remarkably, although *Ptgs2* transcript levels were similarly induced by cytokines alone or in combination with BL001, the secretion of the PTGS2 product PGE_2_ was 5-fold higher in the combined treatment (BL001 and cytokines) as compared to levels in cytokines treated islets (**Figure 4e**). This BL001-mediated and specific increase in PGE_2_ was abolished in *Ptgs2* silenced islets (**Figure 4e**).

**FIGURE 4:**
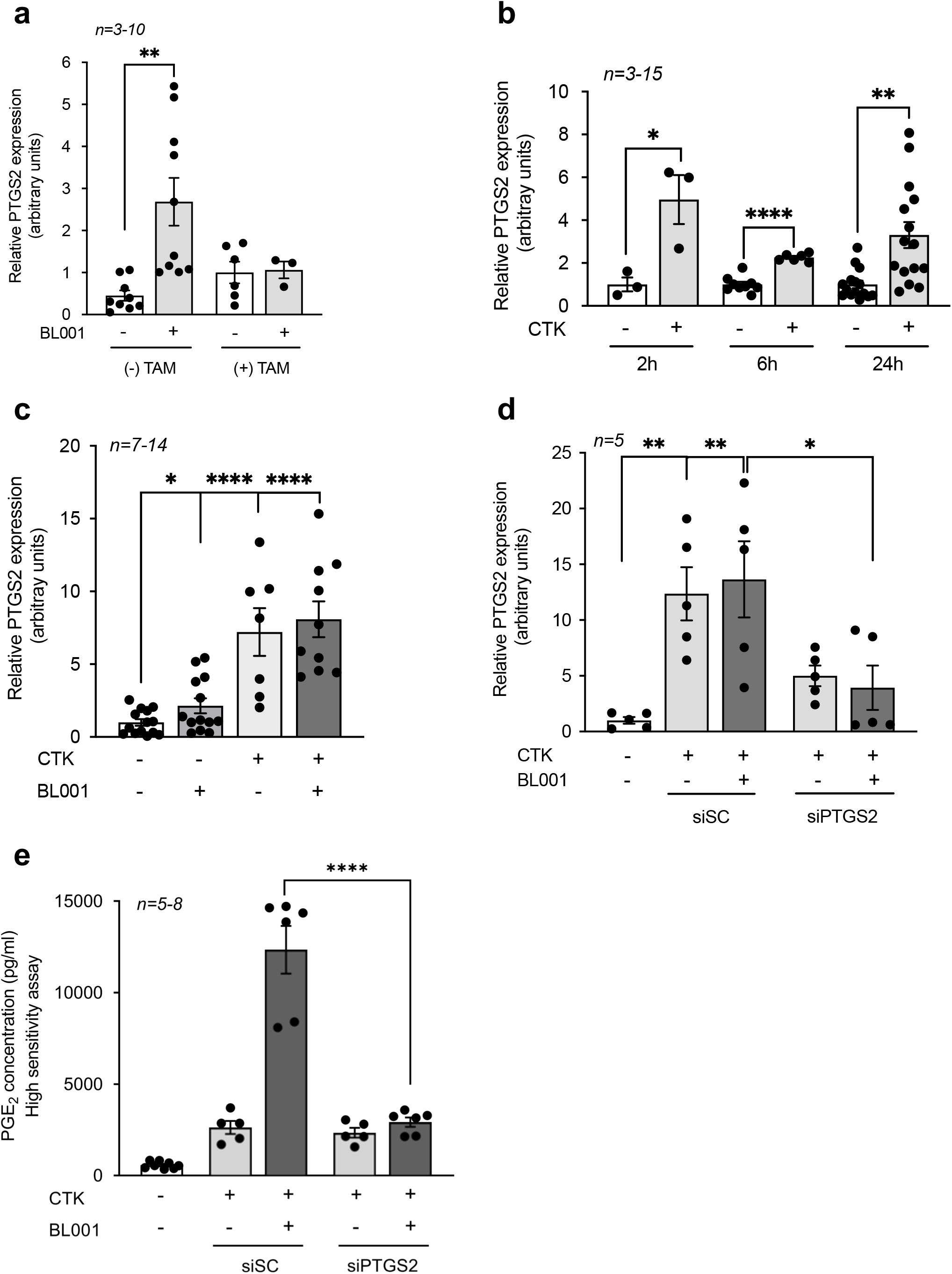
PTGS2 is a downstream target of the BL001/LRH1 signaling pathway. *Ptgs2* expression levels were assessed in (**a**) islets isolated from IndβLRH-1 mice treated or not with TAM, (**b**) islets treated with a cytokine cocktail for 2, 6 and 24 hours, (**c)**islets treated with or without cytokines and BL001 for 24 hours and (**d**) siSC or siPTGS2 transfected islets treated or not with cytokines and BL001. *Ptgs2* expression levels were normalized to either the housekeeping gene *Gapdh* or *Cyclophilin*. (**e**) Secreted PGE_2_ levels were assessed by ELISA in the culture media of siSC and siPTGS2 transfected islets treated or not with cytokines and BL001. n= number of mouse islet preparation. Results are expressed as means + s.e.m. *p < 0.05, **p < 0.01, and ****p < 0.0001 Student’s t test.

### Islets lacking PTGS2 are refractory to BL001-mediated protection against cytokines

We next assessed the impact of silencing PTGS2 on BL001-mediated islet cell survival after cytokines challenge. Hallmarks of cytokine-mediated cell death via the intrinsic apoptotic pathway include the release of cytochrome C from mitochondria resulting in cleavage of PARP ^36^. Consistent with this, cytochrome C release and PARP cleavage were induced/increased in cytokine exposed siSC-treated islets while BL001 rescinded these apoptotic events (**Figure 5a-d**). In contrast, PTGS2 silenced islets were completely refractory to the protective effect of BL001 under cytokine attack, as assessed by the presence of cytochrome C release and increased PARP cleavage (**Figure 5a-d**). In the intrinsic apoptotic pathway, cellular stress leads to the stimulation and translocation of BAX to the mitochondria, an event that permeabilizes the outer membrane, resulting in the release of cytochrome C ^36^. Accordingly, BL001 inhibited *Bax* expression levels in siSC-treated islets while in PTGS2 silenced islets, BL001 treatment failed to repress *Bax* transcript levels, consistent with increased apoptosis (**Figure 5e**).

**FIGURE 5:**
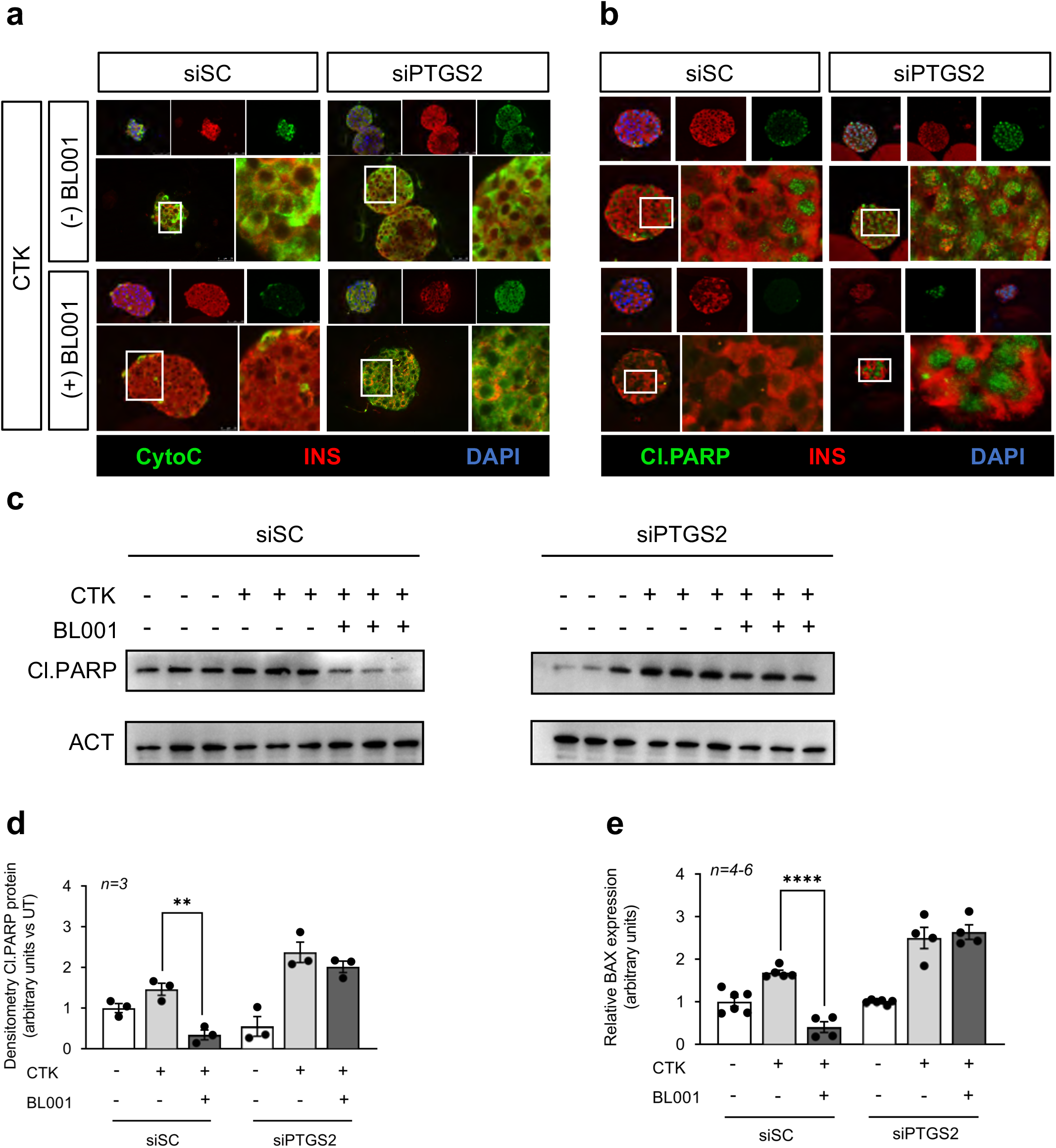
PTGS2 silenced islets are refractory to the protective effect of BL001 under cytokine attack. Representative immunofluorescence images of embedded islets transfected with either a control siRNA (siSC) or siPTGS2 and treated or not with CTK or/and with BL001. Islet slices were stained for either (**a**) Cytochrome C (CytoC, green) or (**b**) cleaved PARP (Cl.PARP, green) along with insulin (INS, red). Nuclei were stained with DAPI (blue). Magnification 40X. Protein levels of cleaved PARP (Cl.PARP) (**c**) and (**d**) quantification thereof were assessed in whole islet cell extracts that were transfected with either a control siRNA (siSC) or siPTGS2 and treated or not with CTK or/and with BL001. Relative protein levels were normalized to ACT. Each line represents an independent experiment. ** p < 0.01 unpaired two-tailed t-test siSC/CTK versus siSC/CTK/BL001 (**e**) Transcript levels of *Bax* were also determined in the same experimental groups. Expression levels were normalized to the housekeeping gene *Cyclophilin*. **** p < 0.0001, unpaired two-tailed t-test siCT/CTK versus siSC/CTK/BL001.

### PTGR1 partially relays the BL001/LRH-1/NR5A2/PTGS2/PGE_2_ signaling axis to islet survival

Several studies indicate that the pro-survival effects of PTGS2 and its product PGE_2_ on islets are conveyed mainly by PTGER4 (*a.k.a*. EP4) while the pro-apoptotic events are transmitted by PTGER3 (*a.k.a*. EP3) for which blockade improves cell survival ^17, 18, 37^. As such we next evaluated whether the islet pro-survival effects of BL001/LRH-1/NR5A2/PTGS2/PGE_2_ were mediated by PTGER4 downstream pro-survival signalling cascade, which include phosphorylation of PKA, CREB and AKT ^38^. Remarkably, only pAKT levels were significantly increased in siSC-treated islets exposed to cytokines and BL001 while pPKA and pCREB were unaffected (**Figure 6a-d**). In contrast, phosphorylation of CREB was increased in cytokines/BL001-treated islets in which PTGS2 was silenced whereas pAKT levels were completely abolished in these islets independent of treatment (**Figure 6a, c and d**). Phosphorylation of PKA was unaffected by silencing and/or treatment (**Figure 6a and b**). As BL001 failed to stimulate both PKA and CREB phosphorylation, we reasoned that PTGR4 may not relay the pro-survival benefits of the LRH-1/NR5A2 agonist. This premise was confirmed as evidenced by the inability of the PTGER4 specific antagonist, L-161,982, to block BL001-mediated islet protection against cytokines (**Figure 6e**).

**FIGURE 6:**
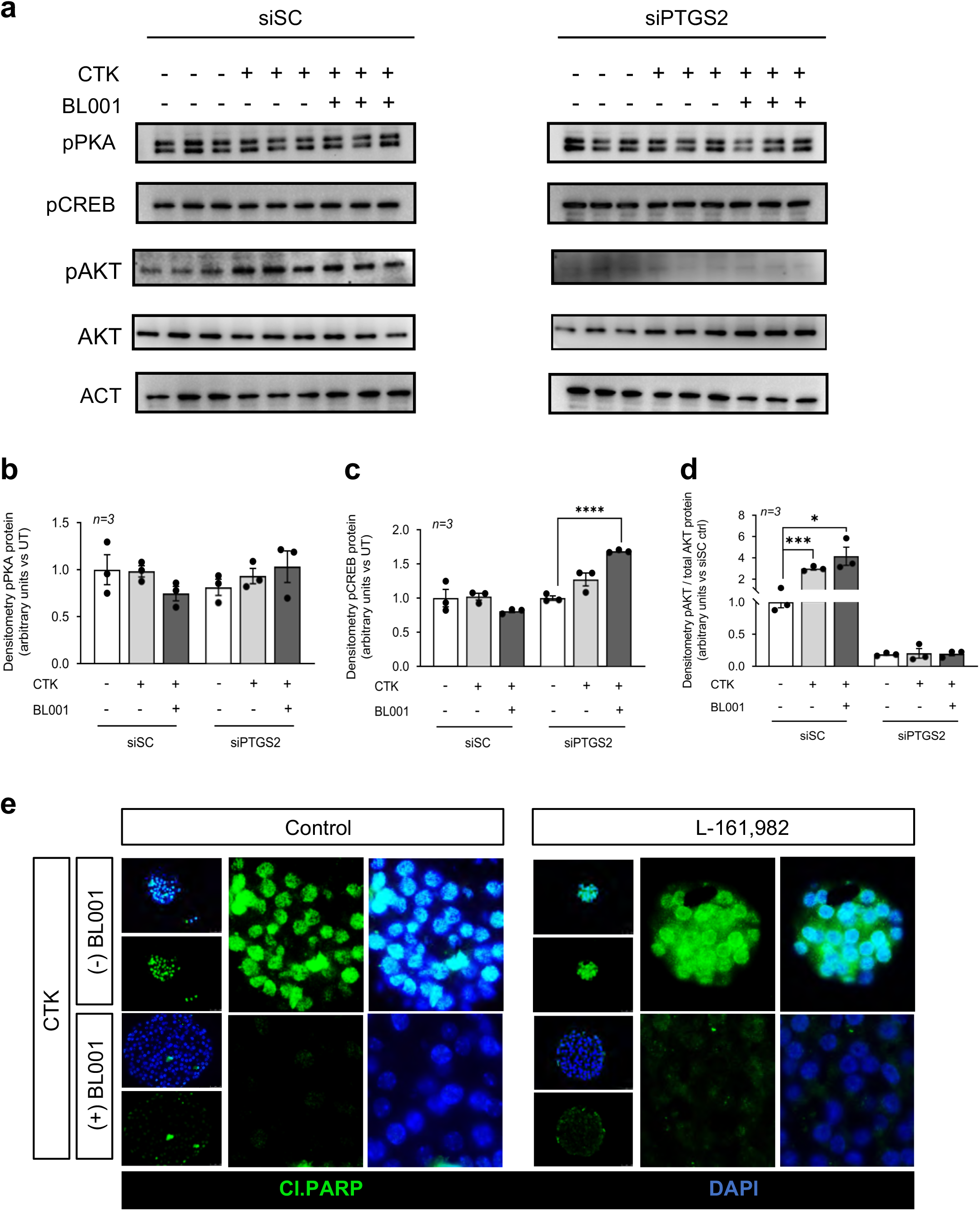
The BL001/LRH-1/NR5A2/PTGS2 anti-apoptotic benefits are not conveyed via PTGER4 signalling. (**a**) Phosphorylated protein levels of PKA (pPKA), CREB (pCREB) and AKT (pAKT) as well as total AKT and quantification thereof (**b**) pPKA, (**c**) pCREB and (**d**) pAKT/AKT were assessed in whole islet cell extracts that were transfected with either a control siRNA (siSC) or siPTGS2 and treated or not with CTK or/and with BL001. Relative protein levels were normalized to ACT for both siCT and siPTGS2. Only 1 representative ACT blot is shown while quantification was performed with matched ACT for each blot. Each lane represents an independent sample. * p < 0.05, *** p < 0.001, **** p < 0.0001 unpaired two-tailed t-test siPTGS2 versus siPTGS2/CTK/BL001. (**e**) Representative immunofluorescence images of embedded islets treated, or not, with a cytokine cocktail, BL001 and/or the PTGER4 antagonist L-161,982. Islet slices were stained for cleaved PARP (Cl.PARP, green) along with DAPI (blue) for nuclei staining. Magnification 40X.

These findings led us to consider whether BL001 could alter the overall expression profile of the 4 PTGER members inducing a shift in PGE_2_ selectivity towards either PTGER1 and 2, also implicated in cell survival. Although not significant, BL001 treated islets displayed an increase in PTGER1 while PTGER2, 3 and 4 were decreased as compared to control islets (**Figure 7a**). Consistent with these tendencies, *in silico* analysis of our previously published transcriptome profile of BL001 treated islets revealed a significantly increase in PTGER1 expression levels as compared to untreated islets (Fold change 1.22, *p*=0.029)^14^. In view of these results, we focused on the potential role of PTGER1 in BL001-mediated cell survival in response to cytokines using the PTGER1 specific inhibitor, ONO-8130. Consistent with our premise, the PTGER1 inhibitor, ONO-8130, blocked the BL001-mediated cell protection against cytokines as gauged by increased cytoplasmic histone-associated DNA fragments and cleaved PARP (**Figure 7b-c**). In contrast, L-161,982 was ineffective in reverting BL001 mediated survival (**Figure 7b**). Taken together these results indicate that PTGR1 is involved in relaying the BL001/LRH-1/NR5A2/PTGS2/PGE_2_ signaling to islet survival.

**FIGURE 7:**
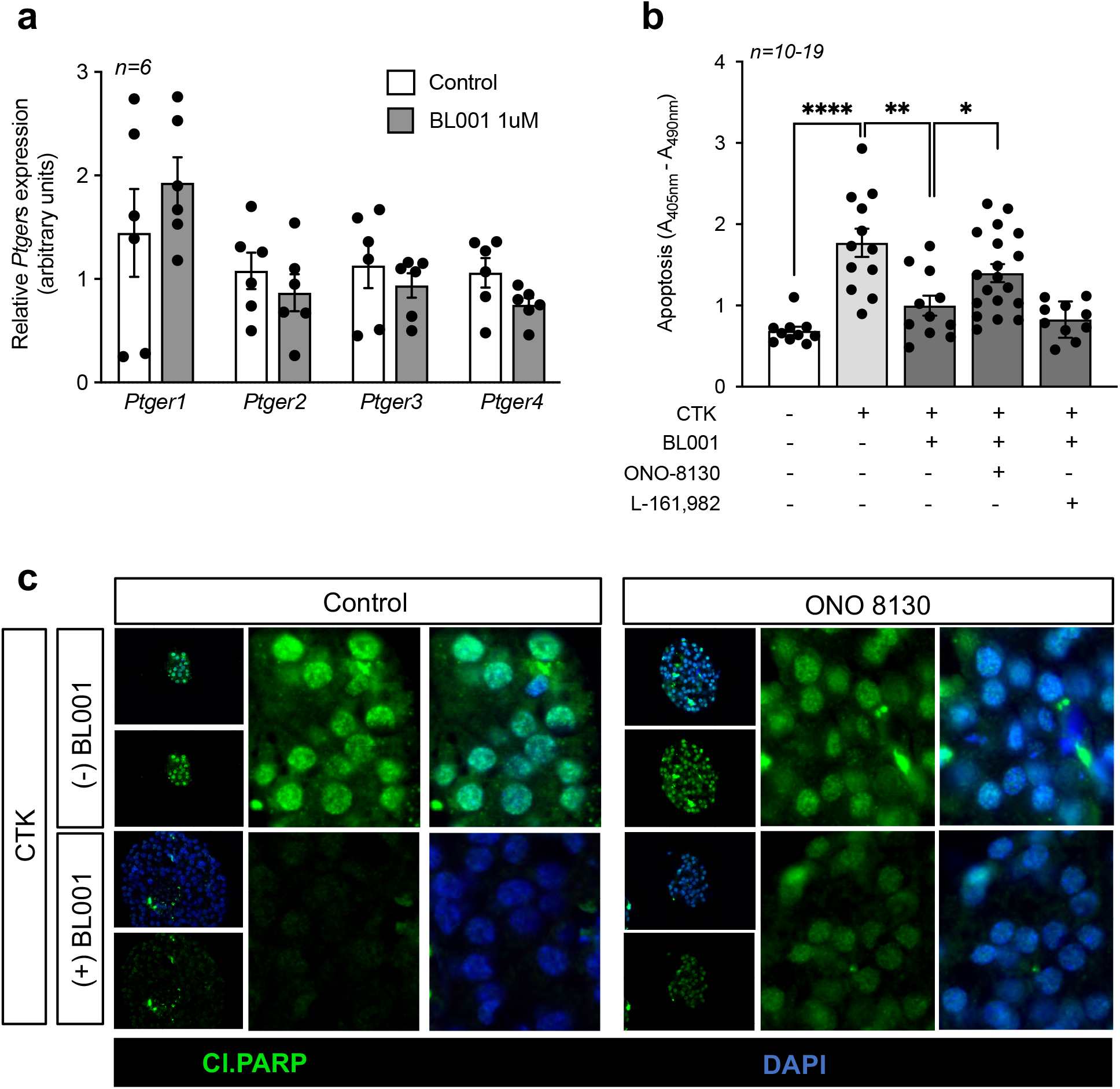
The PGE_2_/PTGER1 signalling pathway mediates the cell protective effect of BL001. (**a**) Expression levels of PTGERs were assessed in islets treated or not with 1uM BL001. Expression levels were normalized to the housekeeping gene *Gapdh.* (**b**) Isolated mouse islets were treated or not with a cocktail of cytokines (CTK), BL001 and antagonists for either PTGER1 (ONO-8130) or PTGER4 (L-161,982). Cell death was assessed by ELISA quantification of mono- and oligo-nucleosomes released by apoptotic cells. ** p < 0.01, unpaired two-tailed t-test CTK versus BL001 and L-161,982 groups. (**c**) Representative immunofluorescence images of embedded islets treated, or not, with a cytokine cocktail, BL001 and/or the PTGER1 antagonist ONO-8130. Islet slices were stained for cleaved PARP (Cl.PARP, green) along with DAPI (blue) for nuclei staining. Magnification 40X.

## DISCUSSION

Herein we disclose that LRH-1/NR5A2 is essential for proper post-natal islet beta cell expansion and that its specific interaction with BL001 regulates the PTGS2-PGE_2_-PTGER1 signalling axis contributing to islet survival and antidiabetic actions of the agonist. These conclusions are based on: 1) the proliferative capacity of post-natal beta cells is severely hampered in the absence of LRH-1/NR5A2, 2) deletion of LRH-1/NR5A2 in adult islets abolishes the pro-survival and anti-diabetic effects of BL001 in STZ-treated mice, 3) silencing of PTGS2 sensitizes BL001-treated islets to cytokine-induce apoptosis and 4) pharmacological blockade of PTGER1 but not PTGR4 partially blocks the anti-apoptotic effects of BL001 in cytokine-treated islets.

A hallmark of neonatal beta cells is their high replicative capacity associated with islet remodelling, organization and maturation which is crucial for subsequent functionality of adult islets in both rodents and humans ^39^. Although our previous studies did not reveal a role of LRH-1/NR5A2 in beta cell replication in adult islets ^7, 14^, we now demonstrate that the nuclear receptor contributes to this process in neonatal islets, and its deletion in beta cells resulted in reduced beta cell number, hypoglycemia and premature death. However, as we detected RIP-Cre activity in the brain we cannot exclude that the severe growth retardation may be, at least in part, consequential to the neuronal deletion of LRH-1/NR5A2, since this nuclear receptor was shown to be implicated in neurogenesis ^40^, a query that is currently under investigation. Interestingly, Dabernat and colleagues reported a comparable phenotype, including hypoglycemia and pre-mature death, in new born mice in which the floxed beta catenin gene (*Catnb^lox/lox^*) was ablated in developing beta cells using the same RIP-Cre transgenic mouse model used herein. They conclude that deletion of beta catenin within the developmental insulin expression domain impedes expansion and subsequent maturation of beta cells, an effect that they argue to be independent of the off-target activity of the RIP promoter detected in the CNS ^41^. Substantiating this premise, repression of TCF7L2 (*a.k.a* TCF4, a nuclear co-activator of beta catenin), in rat newborn pups using antisense morpholinos, blunted beta cell replication resulting in an almost 30% reduction in beta cell mass ^42^. Contextually, LRH-1/NR5A2 cooperates with the beta catenin/TCF7L2 complex to promote cell proliferation through activation of *c-myc*, *Ccne1* and *Ccnd1*^43^. High expression levels of *c-myc* were shown to be correlated with the high replicative capacity of juvenile islets while low levels associated with poor replication in adult islets ^44^. As such, the failure of beta-catenin/TCF7L2 to complex with LRH-1/NR5A2 and downstream activation *c-myc*, as well as *Ccne1* and *Ccnd1* may explain the reduced neonatal beta cell proliferation in ConβLRH-1^-/-^KO mice. Interestingly, in adult islets, we have shown that overexpression of LRH-1/NR5A2 was unable to enhance *Ccne1* and *Ccnd1* expression and beta cell replication, likely due to low levels of beta catenin ^7^. Our findings reinforce the concept that LRH-1/NR5A2 regulatory networks are temporally and cell type restricted through interaction partners that are differentially expressed during development, adulthood and aging.

Although our previous *in vivo* studies established that BL001 treatment blocked/reverted progression of hyperglycemia in 3 independent mouse model of diabetes, the specificity of the agonist for LRH-1/NR5A2 and its MoA remained elusive. We now authenticate the exclusivity of LRH-1/NR5A2 to convey the pro-survival and anti-diabetic therapeutic benefits of BL001 in islets rebutting any potential off-target effects of BL001 via binding to other receptors such as SF-1 (*a.k.a*. NR5A1), the second NR5A family member that share similarities with LRH-1/NR5A2 in both the protein ligand binding domain and DNA binding site ^45^. Of particular importance was the finding that PTGS2 expression was blunted in BL001/TAM-treated IndβLRH-1^-/-^KO mice. As a consequence, cytochrome C release and PARP cleavage were increased upon cytokine insult. These results confirm that PTGS2 is a *bona fide* downstream target of the BL001/LRH-1/NR5A2 anti-apoptotic signalling pathway in islets. Remarkably, *in silico* analysis failed to identify LRH-1/NR5A2 consensus DNA binding sequence (T/C-CAAGG-T/C-CA) within the proximal promoter region of the *Ptgs2* gene indicating that the nuclear receptor does not directly regulate this gene. Notwithstanding, expression levels of several transcription factors including NFκB1 (Fold change 1.23, *p*=0.015), TCF7L2/TCF4 (Fold change 1.36, *p*=0.032), C/EBP (Fold change 1.37, *p*=0.002), ATF4 (Fold change 1.3, *p*=0.029) and ATF5 (Fold change 1.38, *p*=0.005) that bind to and regulate *Ptgs2* transcription were significantly increased in a transcriptomic analysis of BL001 treated islets ^14, 46^. Noteworthy, TCF7L2/TCF4, ATF4 and ATF5 were all implicated in rodent beta cell survival under stress conditions, independent of their regulatory action on the *Ptgs2* gene ^47, 48, 49^. As such we argue that pro-survival signals in response to stress converge onto *Ptgs2* expression in an attempt to re-establish cellular homeostasis and preventing apoptosis, a process that is facilitated by BL001-mediated activation of LRH-1/NR5A2. Such pro-survival function of *Ptgs2* was highlighted in a study in which its deletion sensitized transgenic mice to STZ-induced hyperglycemia ^17^. In the same vein, PGE_2_ supplementation reduced caspase 3 activity in mouse islets ^50^. In contrast, another study reported that sustained *Ptgs2* overexpression in islet beta cells induced hyperglycemia in hemizygous transgenic mice. Notwithstanding, these mice did not exhibit increased apoptosis ^51^. These data argue against a role of PTGS2 in beta cell apoptosis and emphasize the importance of fine-tuning its levels and activity to prevent beta cell dysfunction. The latter is also likely contingent on the expression levels and activity of the 4 PTGERs that dictate the overall impact of PTGS2, as revealed by either blockade of the pro-apoptotic PTGER3 or activation of the anti-apoptotic PTGER4 that increase beta cell survival in the presence of cytokines ^18^.

Our data reveal that BL001 did not augment *Ptgs2* expression above levels induced by cytokines alone while its product, PGE_2_ was robustly increased by the LRH-1/NR5A2 agonist. The latter indicates a post-translational regulation of PTGS2, as reported for the case of inducible nitric oxide synthase that binds, s-nitrosylates, and enhances PTGS2 activity ^52^. Similarly, the nonreceptor Src-family kinase FYN was recently shown to phosphorylate and increase PTGS2 activity resulting in elevated production of PGE_2_ without altering protein steady state levels in prostate cancer cells ^53^. As a consequence of increased PGE_2_, prostate cancer cells were refractory to sanguinarine-induced apoptosis similar to our findings with BL001/cytokine-treated islets ^54^. In parallel, another member of the Src-family of kinases, LYN was also shown to phosphorylate PTGS2, albeit at a different site than FYN, however its impact on PTGS2 activity has not yet being studied ^53^. LYN is regarded as a master suppressor of immune cell activation and its global knockout in mice leads to an autoimmune disorder with similarities to human systemic lupus erythematosus ^55^. In addition to immune cells, LYN is also expressed in pancreatic endocrine cells (https://www.proteinatlas.org/ENSG00000254087-LYN/celltype/pancreas) and, more importantly, its levels are midly but significantly increased by BL001 treatment (Fold change 1.17, *p*=0.008)^14^. Taken together, it is tempting to speculate that BL001-mediated activation of LRH-1/NR5A2 enhances LYN expression in islets, which under stress conditions such as exposure to cytokines will phosphorylate PTGS2 increasing its activity and PGE_2_ production leading to immune modulation and increased cell survival. This hypothesis is currently under investigation.

Downstream signalling of PGE_2_ is conveyed via binding to one or several of the 4 PTGERs with the net cellular physiological outcome, i.e. survival versus death, contingent on the expression levels of these receptors as well as their individual affinity for the prostaglandin in a define environment. For example, expression of the PTGER3 is increased in T2DM patient islets as well as in T2DM mouse models and is associated with beta cell dysfunction and apoptosis ^18^. Blockade of PTGER3 in db/db mice prevents beta cell death ^34, 37^. In contrast activation of PTGER4 enhanced beta cell survival in response to cytokines ^18^ and, as such, we selected this receptor as the ideal candidate to mediate the effect of BL001. This assumption was refuted as BL001 did not increased phosphorylation of PKA and CREB, two downstream target kinases of PTGER4 signaling and the pharmacological inhibition of PTGER4 failed to block BL001-mediated cell survival in response to cytokines. In contrast, AKT phosphorylation, another downstream target of PTGER4, was increased by cytokines alone and in combination with BL001 while silencing of PTGS2 completely abolished phosphorylation independent of treatment indicating that PGE_2_ is a key regulator of AKT phosphorylation levels in beta cells. Nevertheless, AKT phosphorylation can also be conveyed by PKCε via the PI3 kinase dependent pathway and has been associated with both beta cell survival and proliferation ^56, 57, 58^. As PKC is a downstream effector of PTGER1, we argue that AKT phosphorylation could be routed through this receptor reconciliating our finding that BL001-mediated islet survival is blocked by an antagonist of PTGER1. Surprisingly, BL001 in combination with cytokines significantly increased CREB phosphorylation as compared to cytokines alone in PTGS2 silenced islets. The latter likely arise as a consequence of alleviating PGE_2_-induced PTGER3 activity releasing the break on adenylate cyclase activity and enhancing CREB phosphorylation ^34^. These results emphasize the complex cross talk and promiscuity among the various receptors in dictating the overall outcome of PGE_2_ on cell physiology pending environmental inputs. This complex networking is further underlined by findings that PGE_2_ stimulates LRH-1/NR5A2 expression levels indicating a feedforward mechanism between the nuclear receptor and PTGS2 ^3^.

As opposed to PTGER4, pharmacological inhibition of PTGER1 negated BL001-dependent beta cell survival under cytokine attack. In contrast to PTGER2, 3 and 4 that modulate cAMP levels, PTGER1 downstream signalling pathway include increasing intracellular calcium as well as diacyl glycerol (DAG) levels via the Gαq coupling protein leading to the activation of PKC as well as NFκB^59^. Consistent with the PTGER1 signalling cascade being activated by BL001, NFκB1 was significantly increased in a transcriptomic analysis of BL001 treated islets ^14^. Moreover, PKCε activators improve islet survival during isolation as well as function after transplantation in mice ^60, 61^. Further substantiating the implication of PTGER1 in relaying the BL001/LRH-1/NR5A2/PTGS2/PGE_2_ survival signal, GLP-1-mediated activation of PKC was shown to inhibit the intrinsic apoptotic pathway via modulation of BAX while an activated variant of AKT prevented PARP cleavage, two events we report herein ^62, 63^. Oddly, PTGER1 exhibits the lowest affinity for PGE_2_. We reason that high concentrations of PGE_2_ generated in response to BL001 combined with the mild increase in PTGER1 expression with the concomitant decrease in PTGER2, 3 and 4 favors a shift in receptor selectivity towards PTGER1, as observed for high concentrations of prostacyclin that favor interaction with the IP receptor coupled to the Gαq protein ^64^.

In summary, our data refine the cellular mechanism by which LRH-1/NR5A2 is implicated in neonatal beta cell mass establishment through cell replication and define the molecular MoA by which the small chemical agonist BL001 specifically activates LRH-1/NR5A2 and the downstream PTGS2/PGE_2_/PTGER1 signalling axis to foster beta cell survival in response to cytokines. Historically, the role of PTGS2 has been associated with pro-inflammatory response and as such selective inhibitors were developed to treat inflammatory diseases such as osteoarthritis and rheumatoid arthritis ^65^. Notwithstanding, other studies have shown that PTGS2 expression is essential for the resolution phase of the pro-inflammatory response in wound healing raising concerns on the use of such PTGS2 inhibitors for the treatment of inflammatory diseases ^66^. The data herein support this opinion and substantiate our concept recently brought forth that the BL001/LRH-1/NR5A2 axis on one hand protects islet cells against cytokines and on the other hand incites the exit of the perpetual unresolved wound healing of T1DM i.e. continuous pro-inflammatory attack, towards an antiinflammatory and pro-regenerative environment, likely conveyed in part by PGE_2_ secretion and signaling between immune and islet cells ^16^.

## Acknowledgements

We thank Dr. Maria José Quintero and Dr. Paloma Dominguez from the Cytometry and Microscopy Core Facility of CABIMER as well as Irene Diaz Contreras for their excellent technical and analysis support. Special thanks to Dr. Jantje M. Gerdes, Dr. Francisco Real, Dr. Jorge Ferrer and Ms. Irene Millan for their expert advice. We acknowledge the support of the pancreatic islet study group of the Spanish Association of Diabetes.

## Authors contribution

N.C.V., P.I.L, E.M.V., and B.R.G. were involved in research design. N.C.V., P.I.L, E.M.V., E.M., L.L.B., M.G., A.M.M., E.N., V.C., A.C., A.L.G., S.Y.R.Z and F.J.B.S. contributed to conducting experiments. N.C.V., P.I.L, E.M.V., M.G., F.J.B.S. and B.R.G contributed to data analysis. A.R., M.G. and F.M. supplied mouse strains and reagents. E.M.V., N.C.V. and B.R.G. wrote the manuscript. All authors commented on the manuscript. B.R.G. is the guarantor of this work and, as such, has full access to all the data in the study and takes responsibility for the integrity of the data and the accuracy of the data analysis.

## Compliance with ethical standards

### Ethical approval

All experimental mouse procedures were approved by the Institutional Animal Care Committee of the Andalusian Center of Molecular Biology and Regenerative Medicine (CABIMER) and performed according to the Spanish law on animal use RD 53/2013. Animal studies were performed in compliance with the ARRIVE guidelines ^20^.

