## Supplemental material for "LRH-1/NR5A2 regulates the PTGS2-PGE_2_-PTGER1 signalling axis contributing to islet survival and antidiabetic actions of the agonist BL001"

Eugenia Martin Vázquez *et al*.

| ID (Mouse) | Forward primer | Reverse primer |
|---|---|---|
| ***Bax*** | CCCTGTGCACTAAAGTGCCC | CTTCTTCCAGATGGTGAGCG |
| ***Cyclophylin*** | ATGGCAAATGCTGGACCAA | GCCATCCAGCCATTCAGTCT |
| ***Gapdh*** | CACCAACTGCTTAGCCCC | TCTTCTGGGTGGCAGTGATG |
| ***Lrh1/Nr5a2*** | AACGATGTCCCTACTGTCG | CATGCGGTCGGCTCTTAC |
| ***Actb*** | GGACCAGATCCAAAAGGACA | GCTCACCCTTACCTGGAACA |
| ***Rsp9*** | GGAGTCACCCACGGAAGTT | CATGTTCAGCCCGTATTTGC |
| ***Ptgs2*** | GATGCTCTTCCGAGCTGTG | GGATTGGAACAGCAAGGATTT |
| ***Ptger1*** | CAGCACTGGCCCTCTTGG | ATGCCACAGCCAAGCAAAAG |
| ***Ptger2*** | CATCTATGGGGCCTCCTTGC | AGAGAGCTGCAGAATTGACCG |
| ***Ptger3*** | TGTCGGTTGAGCAATGCAAG | CAGCTGGTCACTCCACATCAG |
| ***Ptger4*** | CAGACTGGTCTTCACCGACC | GGAATGGTACCTCCAACCTCA |

**TABLE S1:** List of primers used in this study

| Primary antibodies | Host | Supplier | Catalog number | Use | Dilution |
|---|---|---|---|---|---|
| **PTGS2 Cox2 (D5H5) XP® Rabbit mAb** | Rabbit | Cell Signalling | #12282 | WB/IF | 1:1000/1:150 |
| **Cleaved PARP Recombinant Anti-Cleaved PARP1 antibody [E51]** | Rabbit | Abcam | #ab32064 | WB/IF | 1:1000/1:100 |
| **Phospho-PKA C (Thr197) (D45D3) Rabbit mAb** | Rabbit | Cell Signalling | #5661 | WB | 1:1000 |
| **Phospho-CREB (Ser133) (87G3) Rabbit mAb** | Rabbit | Cell Signalling | #9198 | WB | 1:1000 |
| **GAPDH (14C10) Rabbit mAb** | Rabbit | Cell Signalling | #2118 | WB | 1:10000 |
| **Phospho-Akt (Ser473) (D9E) XP® Rabbit mAb** | Rabbit | Cell Signalling | #4060 | WB | 1:2000 |
| **Akt Antibody** | Rabbit | Cell Signalling | #9272 | WB | 1:1000 |
| **Monoclonal Anti-b-Actin** | Mouse | Sigma-Aldrich | #A5441 | WB | 1:20000 |
| **Anti-GFP antibody** | Goat | Cell Signalling | ab6673 | IF | 1:200 |
| **Monoclonal Anti-Glucagon antibody produced in mouse** | Mouse | Sigma-Aldrich | G2654 | IF | 1:200 |
| **Glucagon Antibody** | Rabbit | Cell Signalling | #2760 | IF | 1:150 |
| **Monoclonal Anti-Insulin antibody produced in mouse** | Mouse | Sigma-Aldrich | #I2018 | IF | 1:300 |
| **Anti-Cytochrome C monoclonal** | Mouse | MBL | #BV-3026-3 | IF | 1:100 |

**TABLE S2:** List of primary antibodies used in this study

| Secondary antibodies | Host | Supplier | Catalog number | Use | Dilution |
|---|---|---|---|---|---|
| **Anti-Mouse IgG (whole molecule)–Peroxidase antibody produced in rabbit** | Rabbit | Sigma-Aldrich | #A9044 | WB | 1:5000 |
| **Anti-Rabbit IgG (whole molecule)–Peroxidase antibody produced in goat** | Goat | Sigma-Aldrich | #A0545 | WB | 1:5000 |
| **IRDye® 680RD Goat anti-Mouse IgG (H + L)** | Goat | LICOR | #926-68070 | WB | 1:15000 |
| **IRDye® 800CW Goat anti-Rabbit IgG (H + L)** | Goat | LICOR | #926-32211 | WB | 1:15000 |
| **Goat anti-Rabbit IgG (H+L) Highly Cross-Adsorbed Secondary Antibody, Alexa Fluor Plus 488** | Goat | Life Technologies | #A32731 | IF | 1:800 |
| **Goat anti-Mouse IgG (H+L) Cross-Adsorbed Secondary Antibody, Alexa Fluor 568** | Goat | Life Technologies | #A11004 | IF | 1:800 |
| **Donkey anti-Mouse IgG (H+L) Highly Cross-Adsorbed Secondary Antibody, Alexa Fluor Plus 555** | Donkey | Life Technologies | #A32773 | IF | 1:800 |
| **Donkey anti-Rabbit IgG (H+L) Highly Cross-Adsorbed Secondary Antibody, Alexa Fluor 647** | Donkey | Life Technologies | #A31573 | IF | 1:800 |
| **Donkey anti-Goat IgG (H+L) Cross-Adsorbed Secondary Antibody, Alexa Fluor 488** | Donkey | Life Technologies | #A11055 | IF | 1:800 |

**TABLE S3:** List of secondary antibodies used in this study

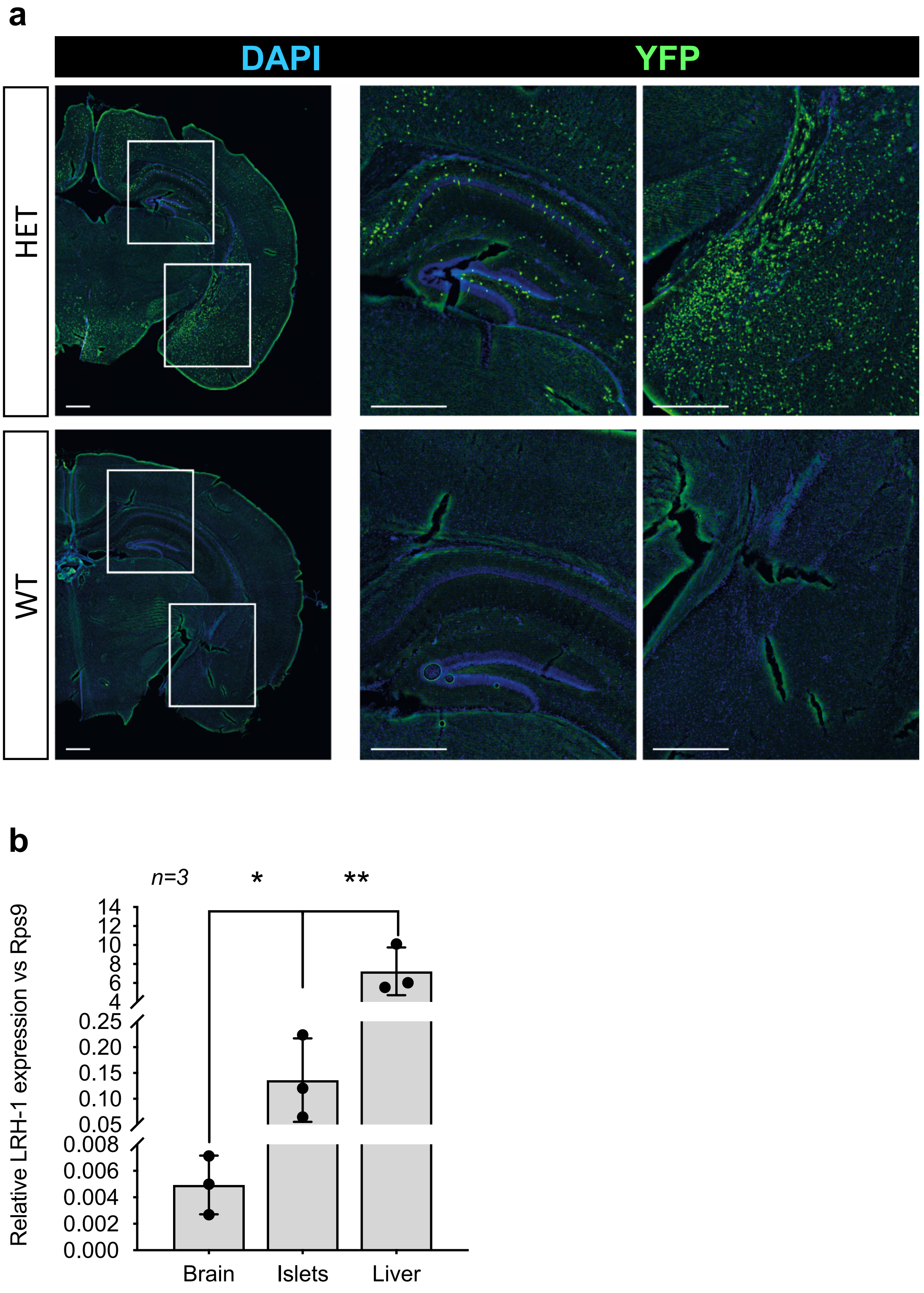


**FIGURE S1: The RIP-Cre is expressed in the brain of IndβLRH-1 mice:** (**a**) YPF immunostaining of central nervous system sections procured from either IndβLRH-1 (HET, upper panel) or LRH-1$^{lox/lox}$::R26-stop-EYFP (WT, lower panel, green). Bar, 0.5 mm (**b**) LRH-1 transcript levels in brain were compared to levels in islets and liver. Relative expression levels were normalized to the housekeeping gene Rps9. *n=3*, independent samples. *p<0.05 and **p<0.001 unpaired t-test Brain versus islets and liver.

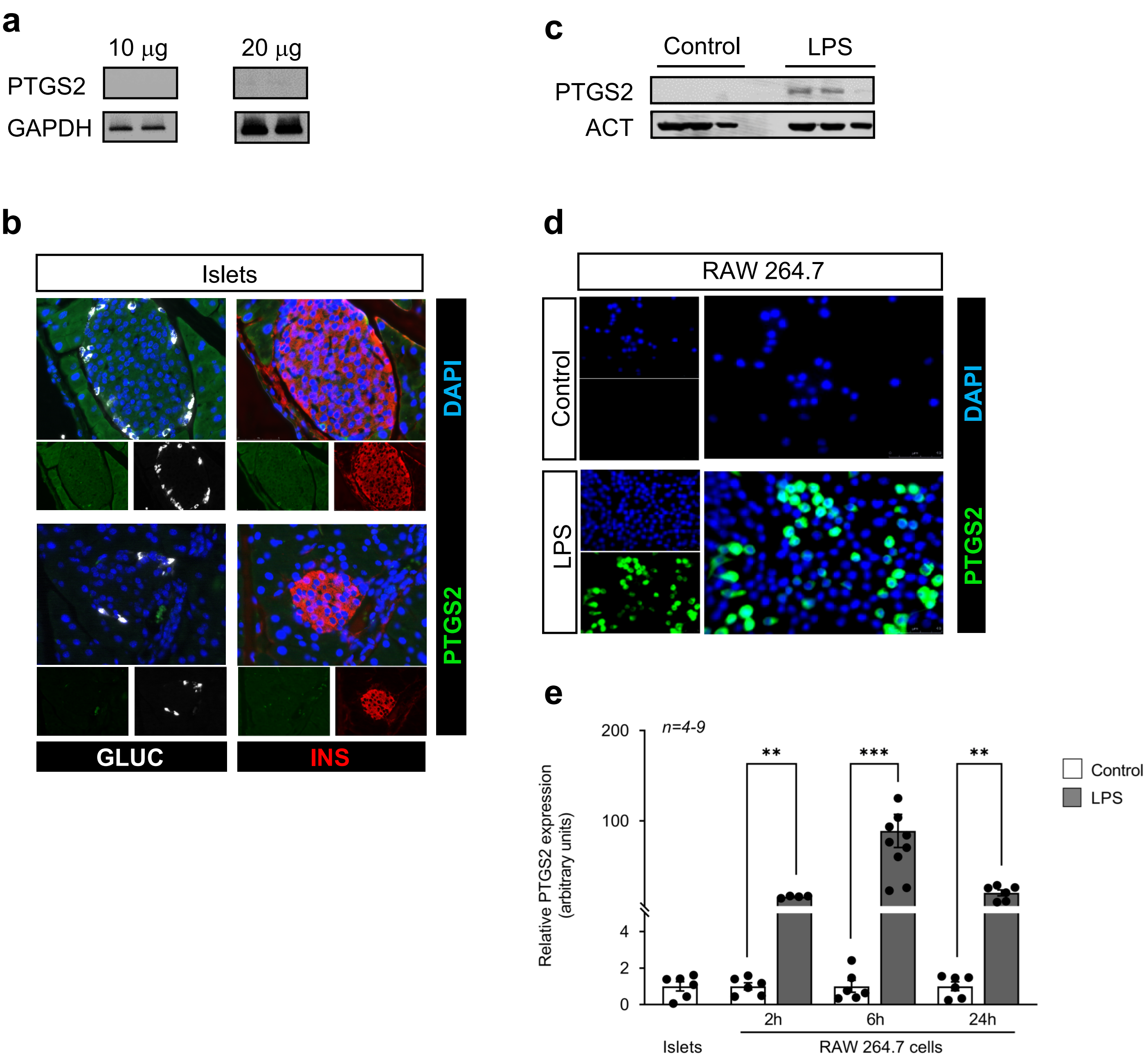


**FIGURE S2. PTGS2 protein expression levels in islets are barely detectable:** (**a**) Western blot analysis of PTGS2 and GAPDH in protein extracts isolated from islets (**b**) Immunostaining of PTGS2 (green), insulin (INS, red) and glucagon (GLUC, red) in pancreas sections. PTGS2 levels in RAW 264.7 cells treated or not with LPS were assessed by (**c**) Western blot and (**d**) immunofluorescence (PTGS2, green and nuclei, blue, Magnification 40X) as well as (**e**) by QT-PCR. Protein levels were normalized to ACTIN while transcript levels normalized to *Gapdh.* Results are expressed as the means + s.e.m. $^{**}p < 0.002$, $^{***}p < 0.001$ Student's t test..
